# ImmunoTyper-SR: A Novel Computational Approach for Genotyping Immunoglobulin Heavy Chain Variable Genes using Short Read Data

**DOI:** 10.1101/2022.01.31.478564

**Authors:** Michael Ford, Ananth Hari, Oscar Rodriguez, Junyan Xu, Justin Lack, Cihan Oguz, Yu Zhang, Sarah Weber, Mary Magglioco, Jason Barnett, Sandhya Xirasagar, Smilee Samuel, Luisa Imberti, Paolo Bonfanti, Andrea Biondi, Clifton L. Dalgard, Stephen Chanock, Lindsey Rosen, Steven Holland, Helen Su, Luigi Notarangelo, NIAID COVID Consortium, Uzi Vishkin, Corey Watson, S. Cenk Sahinalp

**Affiliations:** National Cancer Institute, NIH, Bethesda, MD; National Institute of Allergy and Infectious Diseases, NIH, Bethesda, MD; Department of Electrical Engineering, University of Maryland, College Park, MD; Department of Biochemistry and Molecular Genetics, University of Louisville, KY; Uniformed Services University of the Health Sciences, Bethesda, MD; Diagnostic Department, ASST Spedali Civili di Brescia, Brescia, Italy; University of Milano-Bicocca-Fondazione MBBM, Monza, Italy

## Abstract

Human immunoglobulin heavy chain (IGH) locus on chromosome 14 includes more than 40 functional copies of the variable gene (IGHV), which, together with the joining genes (IGHJ), diversity genes (IGHD), constant genes (IGHC) and immunoglobulin light chains, code for antibodies that identify and neutralize pathogenic invaders as a part of the adaptive immune system. Because of its highly repetitive sequence composition, the IGH locus has been particularly difficult to assemble or genotype through the use of standard short read sequencing technologies. Here we introduce ImmunoTyper-SR, an algorithmic method for genotype and CNV analysis of the germline IGHV genes using Illumina whole genome sequencing (WGS) data. ImmunoTyper-SR is based on a novel combinatorial optimization formulation that aims to minimize the total edit distance between reads and their assigned IGHV alleles from a given database, with constraints on the number and distribution of reads across each called allele. We have validated ImmunoTyper-SR on 12 individuals with Illumina WGS data from the 1000 Genomes Project, whose IGHV allele composition have been studied extensively through the use of long read and targeted sequencing platforms, as well as nine individuals from the NIAID COVID Consortium who have been subjected to WGS twice. We have then applied ImmunoTyper-SR on 585 samples from the NIAID COVID Consortium to investigate associations between distinct IGHV alleles and anti-type I IFN autoantibodies which have been linked to COVID-19 severity.

## 1 Introduction

Recent groundbreaking sequencing technologies producing long reads with low error profiles have allowed researchers to uncover the last hidden secrets of the DNA portion of the human genome. While these novel approaches represent significant steps in high fidelity haplotyping of complex and repetitive regions of the human genome[1], they are expensive and not yet scalable to the population level; furthermore they are reported to have issues with (subclonal) somatic mutations, limiting their potential clinical utility[2].

The vast majority of available genome sequencing data, especially clinical sequence data is, and, at least in the next 3-5 years, will be generated with short read technology. Leveraging the enormous quantity of short read Whole Genome Sequencing (WGS) data available for genomic locus characterization provides an opportunity to gain biological insights into population-level diversity and disease associations. Together, such insights can help inform the development of low-cost clinical genotyping approaches that can be used to guide healthcare decisions. For example, the NIH’s NIAID COVID Consortium^1^ is a large collaborative effort that has generated and analyzed Illumina short read WGS data from COVID-19 patients “with the aim of discovering monogenetic and multigenic variants that contribute to disease susceptibility, severity and treatment outcomes”^2^. Unfortunately the characteristics of short read sequencing technologies make genotyping difficult for repetitive and homogeneous loci across the human genome.

One such challenging but biologically significant region is the immunoglobulin heavy chain (*IGH*) locus on chromosome 14. The *IGH* locus harbors the variable (*IGHV*), diversity (*IGHD*), joining (*IGHJ*), and constant (*IGHC*) genes (or gene segments) that encode the building blocks of expressed B cell receptors (BCRs) and antibodies (Abs). The variable domain of BCRs and Abs is composed of *IGHV, IGHD*, and *IGHJ* gene segments, and is responsible for engaging directly with antigen, playing a critical role in identifying and neutralizing pathogenic viruses and bacteria. Despite the importance of BCRs and Abs in the adaptive immune system, our understanding of population-level genetic diversity in the *IGH* locus and its contribution to Ab function in disease remains limited[3, 4, 5]. The severe impact of the COVID-19 global pandemic stresses the need for the development of tools and approaches to more fully characterize critical immune genes, especially those involved in the development of critical neutralizing Abs. Unfortunately, due to its intricate sequence composition, *IGH* has remained stubbornly resistant to large scale, high resolution characterization.

*IGH* is one of the most complex and dynamic regions of the human genome[3] - it is known to contain many large structural variants (SVs), including segmental duplications, large insertions and deletions, and other copy number variants (CNVs)[6, 7, 8]. Of the *IGH* gene segments, *IGHV* is the most extensive gene group, and the ∼ 800 Kb - 1 Mb region of *IGH* in which this gene family resides has proven particularly challenging to genotype using short read WGS data, due to its repetitive and homogeneous nature[9, 10]. A given individual is likely to have upwards of 50 functional and ORF *IGHV* gene copies, and at least twice as many non-functional pseudogene copies, collectively residing across the primary *IGH* locus on chromosome 14 and two *orphon loci* located on chromosomes 15 and 16[8, 6]. These gene segments are short - between 165 bp and 305 bp (with a mean of 291 bp) - and are highly similar to others in terms of sequence composition, with 40% of the known functional alleles having a *sequence similarity* of *>* 98% with at least one other allele in a different gene^3^.

While germline *IGH* genotypes have generally been overlooked in GWAS and other disease association studies, partially due to genotyping challenges, there is increasing interest in investigating the effect of *IGH* germline genetic variation on the adaptive immune system and disease[3, 5]. In fact *IGHV* germline genetic variation has recently been associated with clinical phenotypes including infectious disease and vaccine response [11, 12, 13, 14], autoimmune/inflammatory conditions[15, 16, 17], and cancer[18].

Of particular interest is COVID-19, which is primarily modulated by the innate immune system[19]. However, there is increasing evidence for an association between anti-type I interferon (IFN) autoantibodies and COVID-19 severity[20, 21, 22]. This presents a potential biological mechanism for an association between *IGHV* germline genetic variation and COVID-19 disease severity, driven by *IGHV* germline alleles coding anti-type I IFN antibodies.

To date, there has been only one published method/computational pipeline for germline *IGHV* genotyping using short read WGS data[23, 24]. Due to the genotyping challenges encountered with *IGHV*, the authors intended their pipeline to be applied on a population level, stating that it “is not intended to be used to accurately genotype individual genomes”. The authors indeed applied their pipeline to a cohort of 109 individuals with the purpose of compiling aggregate measures of variation, and performed gene-level genotyping, with allele-level granularity on 11 (of the 56) *IGHV* genes.

One other previously published method, ImmunoTyper[25], performs *IGHV* genotype and copy number analysis using standard single molecule real-time (SMRT) Pacific Biosciences (PacBio) long read sequence data. As a result, its application has been limited by the paucity of publicly available long read WGS datasets. ImmunoTyper can not be easily modified to handle short read WGS data because it relies on extracting full length *IGHV* segments from long PacBio reads (which are then clustered using a facility location formulation, similar to balanced k-means clustering). Short reads, on the other hand, only partially cover *IGHV* segments.

More recently Rodriguez et al.[8] have published a novel technique called IGenotyper, which can haplotype the entire *IGH* region at high resolution using targeted ultra-deep *long read sequencing* and *de novo* assembly. This represents a critical step towards characterizing *IGH* haplotype diversity and heterogeneity, however since it requires a custom sequencing approach, it is expensive to scale and as a result is more suitable for high-resolution haplotype characterization rather than high-throughput *IGHV* genotyping.

There are several published methods for inferring germline *IGHV* genotypes using adaptive immune receptor repertoire sequencing (AIRR-seq) data as input[26, 27]. This approach benefits from not having to contend with pseudogene sequence, however it introduces noise that arises from somatic hypermutation. Furthermore, these methods can only provide calls for germline *IGHV* alleles that are expressed in the B cell population at the time of sampling, and thus are susceptible to missing *IGHV* alleles with lower expression.

In this paper we introduce ImmunoTyper-SR, a novel computational approach for genotype and CNV analysis of functional germline *IGHV* genes using short read WGS data, using a database of known *IGHV* sequences as a reference. ImmunoTyper-SR is based on a novel combinatorial optimization formulation that aims to minimize the total edit distance between the reads and their assigned alleles while maintaining additional constraints on the number and distribution of reads across each allele identified. This approach significantly extends our previous work on long read based *IGHV* genotyping by addressing additional challenges introduced through the use of short reads. As a result, ImmunoTyper-SR is the first short read based germline *IGHV* genotyping tool that offers allele-level resolution.

We have validated ImmunoTyper-SR on 12 individuals with diverse genetic backgrounds from the 1000 Genomes Project[28] (1kGP), by comparing ImmunoTyper-SR genotype calls on Illumina WGS data from the NYGC[29] against targeted long read-based *IGH* assemblies generated using IGenotyper[8]. We then applied ImmunoTyper-SR to WGS data from a cohort of 585 individuals from the NIAID COVID Consortium (from here on the “NIAID cohort”) to investigate associations between *IGHV* genotypes, type I IFN autoantibodies and COVID-19 disease severity. The cohort includes nine individuals who have been independently sequenced twice; this subcohort provides additional means to assess the robustness of ImmunoTyper-SR on clinical sequencing data. We observed that ImmunoTyper-SR is able to produce *IGHV* accurate allele and CNV calls on both data sets, demonstrating its feasibility on Illumina WGS data with read lengths of 150 bp and moderate genome coverage. We finally employed a permutation test on the alleles identified by ImmunoTyper-SR to determine those strongly associated with anti-type I IFN autoantibody activity - within the limitations of the size and demographic composition of the NIAID cohort.

## 2 Methods

### 2.1 WGS Read Recruitment

A mapped WGS sample is provided to ImmunoTyper-SR in the BAM file format. Reads are extracted if they share any mapping to *IGHV*, the *IGHV* orphons on chromosome 15 and 16, or any other loci that share sequences similar to *IGHV*.

### 2.2 Allele Candidate Assignment

The extracted reads are then mapped to a database of *IGHV* allele reference sequences using BWA-MEM[30], with the `-a` option. The allele reference database was created by combining the current IMGT allele database[31] (including pseudogene and orphon alleles) with additional germline alleles (currently unpublished), resolved by Rodriguez et al. using IGenotyper[8]. While these novel alleles are as-yet unpublished, they are resolved using PacBio sequencing on an independent dataset that is not subject to the noise present in the 1kGP samples (see Section 2.4 *Annotation of 1kGP Sample Assemblies*).

A read is putatively assigned to a single candidate allele or set of candidate alleles if it has a mapping of at least 50 bp to the allele reference sequence. A read can have a truncated mapping to an *IGHV* allele if and only if the truncation occurs either at the beginning or the end of the allele, such that truncated mappings that do not start on the first base of the allele, or end on the last base, are removed.

The edit distance between a given read and candidate allele is taken from the `NM` tag of the mapping, which is used below.

### 2.3 Allele Assignment

Reads are assigned to one of their candidate alleles through an Integer Linear Programming (ILP) approach, which aims to minimize the total sequence variation (i.e. edit distance) between assigned reads and alleles, while matching the sequence depth and variance of the given WGS sample. The ILP is solved using the Gurobi package[32].

As previously noted by Luo et al.[23, 24], a correct read assignment for a given allele will have a read depth in the shape of a trapezoid, as the number of reads having the minimum 50 bp of mapped sequence will decrease towards the ends of the allele reference sequence. This problem could be avoided by including the non-coding flanking sequence in Section 2.3, however this technique may be complicated by the well known lack of characterized intergenic *IGH* sequences and haplotypes. As a result, we have opted to omit the flanking sequences in order to minimize any chances of genotype biases caused by reference haplotype sequences.

#### 2.3.1 Sampling Sequence Depth and Variance

To ensure that our read assignment matches the underlying sample’s sequencing characteristics, we empirically calculate the sequencing depth and variance from each sample by examining the WGS mapping at a representative locus. We use exon 327 of the TTN gene, located on chromosome 2. This region was selected for being one of the longest exons in the genome, and thus provides a ∼ 17 kbp sampling region that is likely to be relatively stable[33]. We further confirmed that there are no other loci in the genome with a similar sequence by simulating 200X 150 bp reads from the exon using ART[34] and mapping them back to GRCh38 using BWA-MEM[30] with the `-a` parameter; indeed, all reads mapped back to the exon region.

The sequencing depth and variance are calculated as the mean and variance of the mapped read depth across exon 327. To obtain the expected sequencing depth variance over the ‘sloped’ regions of the trapezoid shape found in a valid *IGHV* read assignment as defined in Section 2.3, we calculate the read depth variance for each position of window of size *read_length* − 50 across all bases of exon 327.

#### 2.3.2 Constraining Coverage to Match Sampled Values

In order to match the assigned read depth and variance to the TTN-sampled values, we must monitor the read depth for each possible assignment that arises during the optimization process defined below. We only monitor the read depth at equally-spaced ‘landmark’ positions, which are chosen for every candidate allele in advance, to reduce optimization time. We can then constrain the mean assigned read depth across landmark positions to be within one standard deviation of the expected read depth using the sequencing depth and variance values calculated in Section 2.3.1. However, this constraint may allow for a read assignment with outlier read depth values at consecutive landmark positions that retain a mean coverage within the bounds. To combat this problem, we group landmarks (in a round-robin process) and set the constraint defined above for each landmark group independently (see Figure 1). This reduces the likelihood that two such balancing outlier landmarks are in the same group.

**Figure 1:**
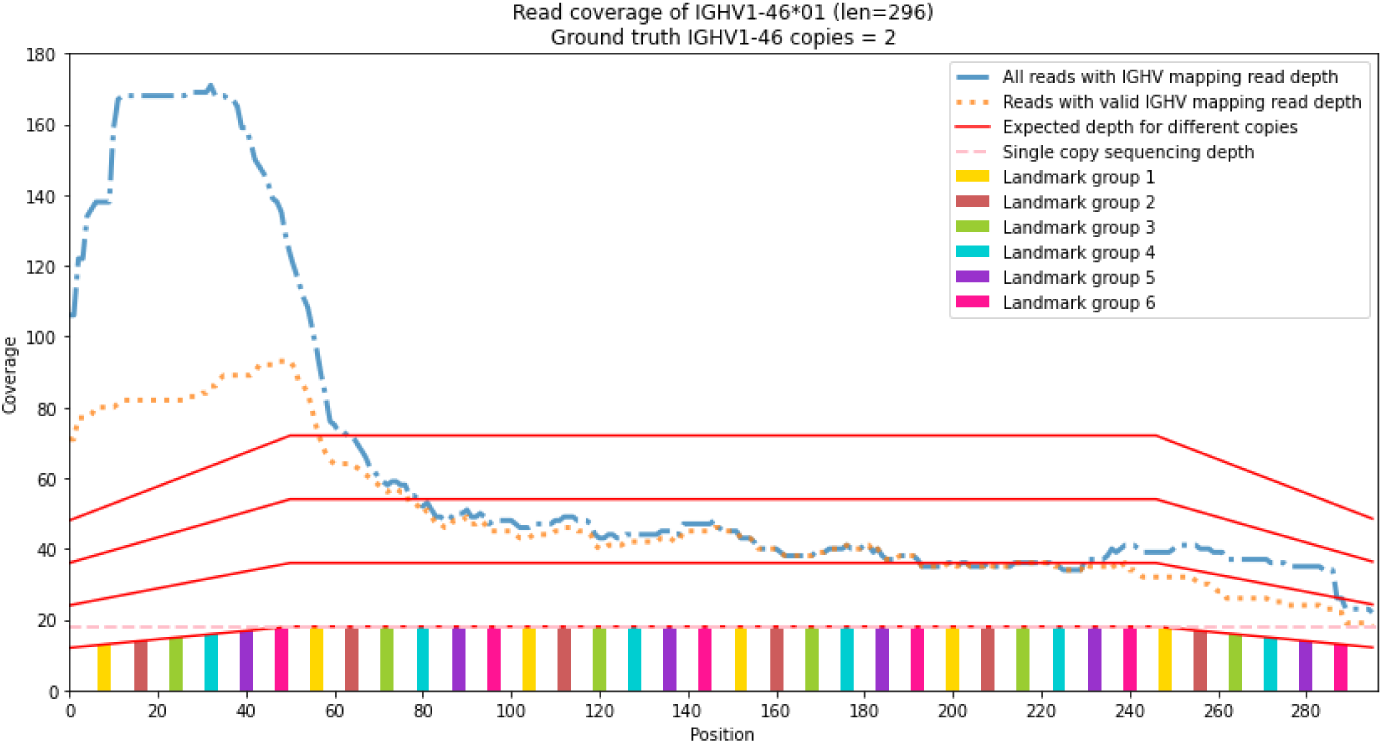
Example of mapped read coverage prior to read ILP model assignment using *IGHV1-46*01* in sample NA19240. As there are two copies in the ground truth, the expected read depth is the second red trapezoidal line from the bottom. Note the mapping and filtered mapping lines (blue and orange, difference explained in Section 2.2) are close to the expected read depth line in the second half of the sequence, but in the first half there are significantly more mappings than expected. The goal of ImmunoTyper-SR is to filter the erroneously mapped reads to match the expected read depth for the ground truth number of copies.

#### 2.3.3 Discarding reads

The read assignment optimization strategy allows for reads to be discarded, to contend with reads that come from non-*IGH* loci which do not have their source sequence characterized in the IMGT allele database, but nonetheless have a mapping due to their similar sequence content. A given read can be discarded with a penalty equal to the expected number of errors for the read, multiplied by a constant factor, which is set to 2, by default. This implies that to be discarded, a read needs twice as many errors as expected relative to to the closest matching allele in the database which allows for a small number of novel variants to exist without the read being discarded.

#### 2.3.4 ILP Model Formulation

##### Definitions

Let 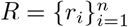 be the set of all WGS reads in the sample.

Let 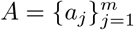 be the set of database alleles. Let **C**_*i*_ be the set of candidate alleles of read *r*_*i*_.

Let **L**_*j,g*_ be a set of landmark nucleotide positions in *a*_*j*_ that are part of landmark group *g*, where *g* is a subset of all positions in *a*_*j*_. **L**_*j*_ is the set of all landmark positions in allele *a*_*j*_.

Let *covers*(*r*_*i*_, *a*_*j*_, *l*) be the set of reads that have *a*_*j*_ as candidate/mapping and cover landmark *l*.

Let *µ*_*l,j*_ and *σ*_*l,j*_ be the expected mean and standard deviation (*SD*) in read coverage of allele *a*_*j*_ for a given landmark *l* ∈ **L**_*j*_ for a single copy of the allele. This is computed by sampling read coverage within

TTN gene’s exon 327 and dividing by two to account for the diploid nature of the exon to estimate empirical mean and SD for the read coverage of landmark *l*.

Let *λ* = user-provided upper bound on the number of standard deviations allowed on the deviation of mean read coverage of any reference allele. Default 1.5.

Let *min_cov* = user-defined proportion of the estimated mean read coverage of any landmark position that the data needs to satisfy. Default 0.3.

Let *e*(*r*_*i*_, *a*_*j*_) = edit distance for the best mapping/alignment of *r*_*i*_ on *a*_*j*_.

Let *expected_errors*(*r*_*i*_) = sequencing error rate × alignment length of primary mapping for *r*_*i*_.

Let *discard_penalty_multiplier* = user-provided penalty for discarding a read. Default 2.

##### Variables

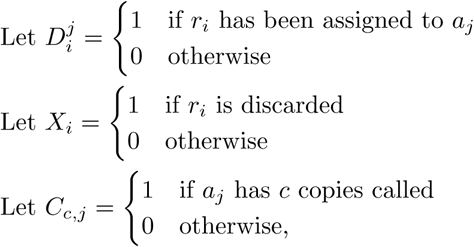

where *c* ∈ *{*0, …, *K}* where *K* is the maximum allowed number of copies per allele.

##### Constraints

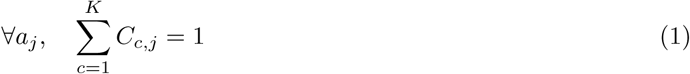

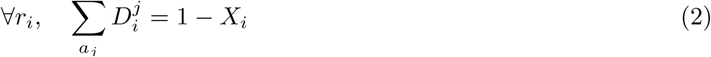

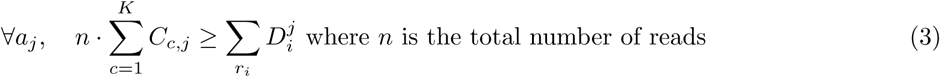

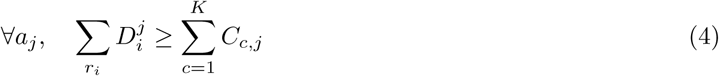

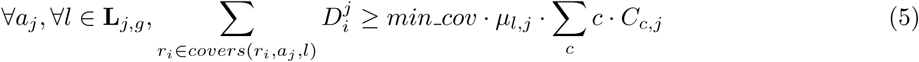

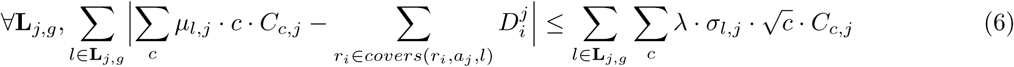

##### Explanations

1. One-hot encoding of all possible copy numbers for each allele.
2. Prevent discarded reads from being assigned to any allele and enforce assignment to at most one copy.
3. If at least one of the reads is assigned to allele *a*_*j*_, then *at least* one copy must be called.
4. If *C*_*j*_ == *c*(*>* 0) (allele *a*_*j*_ is called and has *c* copies), there must be *at least c* reads assigned to *a*_*j*_, to ensure there is no allele copy with zero reads assigned.
5. Each landmark position in **L**_*j,g*_ should have a minimum read coverage proportional to the expected coverage, if allele *a*_*j*_ is called.
6. If *c* copies of allele *a*_*j*_ are called, the deviation of read coverage of a group of landmark positions away from the estimated mean is bounded.

##### Objective Function

###### Minimize

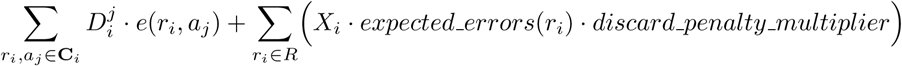

### 2.4 Annotation of 1kGP Sample Assemblies

We created *IGHV* gene and pseudogene annotations for each haplotype from the 12 1kGP *IGH* assemblies to act as ground truth against which to compare the ImmunoTyper-SR allele calls. The complete allele database was mapped against each assembly using Bowtie2[35] (with `-af --end-to-end --very-fast` parameters). The mapping results are then grouped by target location, and the best mapped allele for each target location is assigned to that gene. Any assignment with an edit distance *>* 1 is manually reviewed.

It is possible for an assembly to have *IGHV* gene copies that have significantly divergent sequence from any allele in the database. This can happen either through a true novel allele, as a result of a sequencing or assembly error, or due to large structural alterations present in the sample haplotype. This latter case is of particular concern as the 1kGP samples employed in this study are derived from lymphoblastoid B cell lines (LCL), which have undergone some degree of V(D)J rearrangement. This can introduce noise in the sequencing dataset, and can result in somatic deletions in a haplotype that may affect one or more *IGHV* genes.

To identify any significantly divergent *IGHV* genes, we generated 150 bp error-free in-silico reads from all sequences in the allele database and mapped them to the haplotypes using BWA-MEM[30] (`-a` parameter). Any contiguous target region not identified above was extracted and mapped back to the allele database using BWA-MEM[30] (`-a` parameter) to identify the most similar allele and edit distance.

### 2.5 Statistical Association between *IGHV* Genotype and anti-Type I IFN Autoantibody Presence

First, the 585 samples from the NIAID cohort were filtered for the presence of *α, β, ω* IFN autoantibody labels. Any label with a “Yes, partially” value was modified to “Yes”, and we combined the individual *α, β, ω* labels into a single binary label indicating presence of any one of the autoantibodies. *IGHV* alleles that were present in at least one of sample’s genotype were chosen as the independent variable, in the form of binary presence/absence labels. A candidate selection logistic regression model was then performed between all such *IGHV* alleles and anti-type I IFN autoantibody presence, and those alleles with a significant *p*-value were then selected as top candidates.

The candidate alleles were applied in a separate logistic regression model to determine the effect size. For significance and multiple test correction, we performed a permutation test, by applying a logistic regression model to the dataset 100,000 times, where the autoantibody labels were randomly shuffled in each iteration. The resulting *p*-value was calculated for each allele as the proportion of the shuffled models that had a more extreme *p*-value than that in the original, unshuffled model.

All logistic regression was performed using the `glm` function in R with the `family=“binomial”` parameter.

### 2.6 Comparison With Luo et al. [24] Pipeline

We implemented the pipeline as described in “Worldwide genetic variation of the *IGHV* and *TRBV* immune receptor gene families in humans”[24]. *IGHV* gene calls and CNVs were compared against *IGH* assembly annotations as described in Section 2.4.

Luo et al. limit allele calling to the following 11 genes that they determine are two-copy genes: *IGHV1-18, IGHV1-24, IGHV1-45, IGHV1-58, IGHV2-26, IGHV3-20, IGHV3-72, IGHV3-73, IGHV3-74, IGHV5-51*, and *IGHV6-1*. Since not all these genes are present in two copies in the 1kGP samples, we further limited allele calls only to those genes that are present in the above list, and for which the pipeline gave CNV calls of two.

As a result, when comparing the allele calls against a sample’s ground truth and ImmunoTyper-SR, we only used those genes that met the pipeline’s allele-calling criteria in *IGHV* assembly.

### 2.7 Measuring Concordance between Doubly-sequenced Samples

To calculate genotype call concordance between two WGS sequences from the same subject of the NIAID subcohort that has been doubly-sequenced, we used the weighted Jaccard similarity coefficient (*J*_*w*_), defined as follows:

Let *s* be a subject who was sequenced twice and *g*_*s,i*_ be the genotype vector corresponding to *i*-th WGS sample of the subject (*i* ∈ *{*1, 2*}*), where *k*-th element of the vector (*g*_*s,i*_[*k*]) is the number of copies of allele *A*_*k*_, as called by ImmunoTyper-SR. Then, the weighted Jaccard similarity coefficient for the subject *s* is:

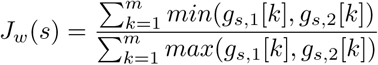

## 3 Results

A primary challenge in germline *IGHV* genotyping is the validation of computationally identified alleles. To date there has been only one *IGHV* genotyping protocol published that uses short read data[23, 24]. A recently published novel *IGH* haplotyping tool[8] has opened the door for high quality *IGH* assemblies which can be used for validation.

*IGHV* genotyping through inference using AIRR-seq is common-place and well established[26, 27], however, we were unable to find any publicly available paired AIRR-seq and WGS datasets that would be suitable for comparison.

### 3.1 Validation using 1kGP Samples

We ran ImmunoTyper-SR on 12 publicly available high-coverage WGS of 1kGP samples, sequenced at ∼ 30X on the Illumina NovaSeq 6000 Platform[29]. These samples have had independent *de novo IGH* haplotyping performed by Rodriguez et al. (unpublished) using a targeted long read sequencing and assembly protocol called IGenotyper[8] tailor-made for the *IGH* region. While this technique represents the most accurate *IGH* haplotyping tool published to date, it is likely that these assemblies are not 100% accurate, as the samples are derived from LCL cell lines. It has been noted that the usage of LCL cell lines can impact *IGH* genotype accuracy due to the presence of somatic V(D)J rearrangement[36]. This may result in somatic deletions or inconsistent read coverage, which can affect the assembly quality, and ultimately ImmunoTyper-SR’s genotype accuracy.

Genotype call results (including the number of copies of each allele) are shown in Figure 2; as can be seen, the mean precision and recall values are respectively 83.7% and 80% for identification of each allele sequence and its copy number exactly. While ImmunoTyper-SR generally provided accurate genotype calls, the range of precision and recall values were significant, with a minimum and maximum F-score of 63% and 88% respectively. These figures improve substantially if some limited noise can be tolerated in the sequence composition or copy number of the allele calls, as discussed in the remainder of the section.

**Figure 2:**
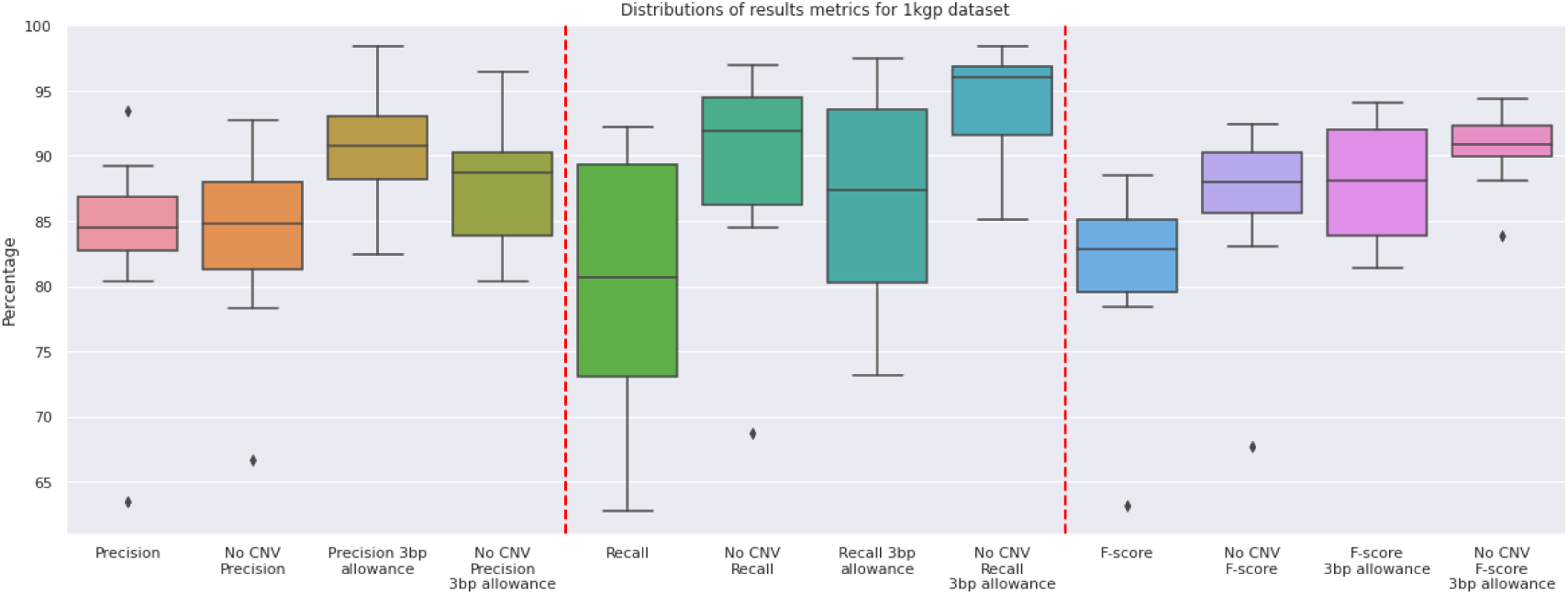
ImmunoTyper-SR’s functional allele call accuracy on 1kGP dataset. No CNV indicates measures accuracy for presence/absence of alleles. 3 bp allowance counts a false positive as a true positive if there is a false negative allele within 3 bp edit distance.

#### 3.1.1 Accuracy is Significantly Impacted by Novel Alleles

As ImmunoTyper-SR is designed to find the closest allele in the database and not call novel alleles and variants, we investigated the relationship between the presence of putative novel variants as a driver of the variation seen in genotype accuracy. It is important to distinguish these putative novel alleles, which are derived from the 1kGP sample *IGH* assemblies, from those validated novel alleles that were added to the allele database as described in Section 2.2 *Allele Candidate Assignment*, which were sourced from an independent, high-quality dataset.

As shown in Figure 4, there is significant correlation between the proportional divergence of a given sample’s *IGHV* alleles from the allele database and ImmunoTyper-SR’s genotype call accuracy.

As previously mentioned, it is likely that some of these novel variants are in fact due to sequencing or assembly errors, or noise from *IGH* rearrangement due to the LCL-sourced samples. Since it is impossible to determine which of the putative novel alleles are valid, we chose not to add these novel sequences to the allele database for fear of increasing noise and model complexity.

#### 3.1.2 High Genotype Accuracy for Distinct Genes

To investigate the genotype accuracy on a gene-by-gene basis, we grouped allele call results by genes across all samples. Figure 3 demonstrates how genes that have high sequence similarity to many alleles in the database (regardless of gene “source” of each such allele) have lower allele call accuracy.

**Figure 3:**
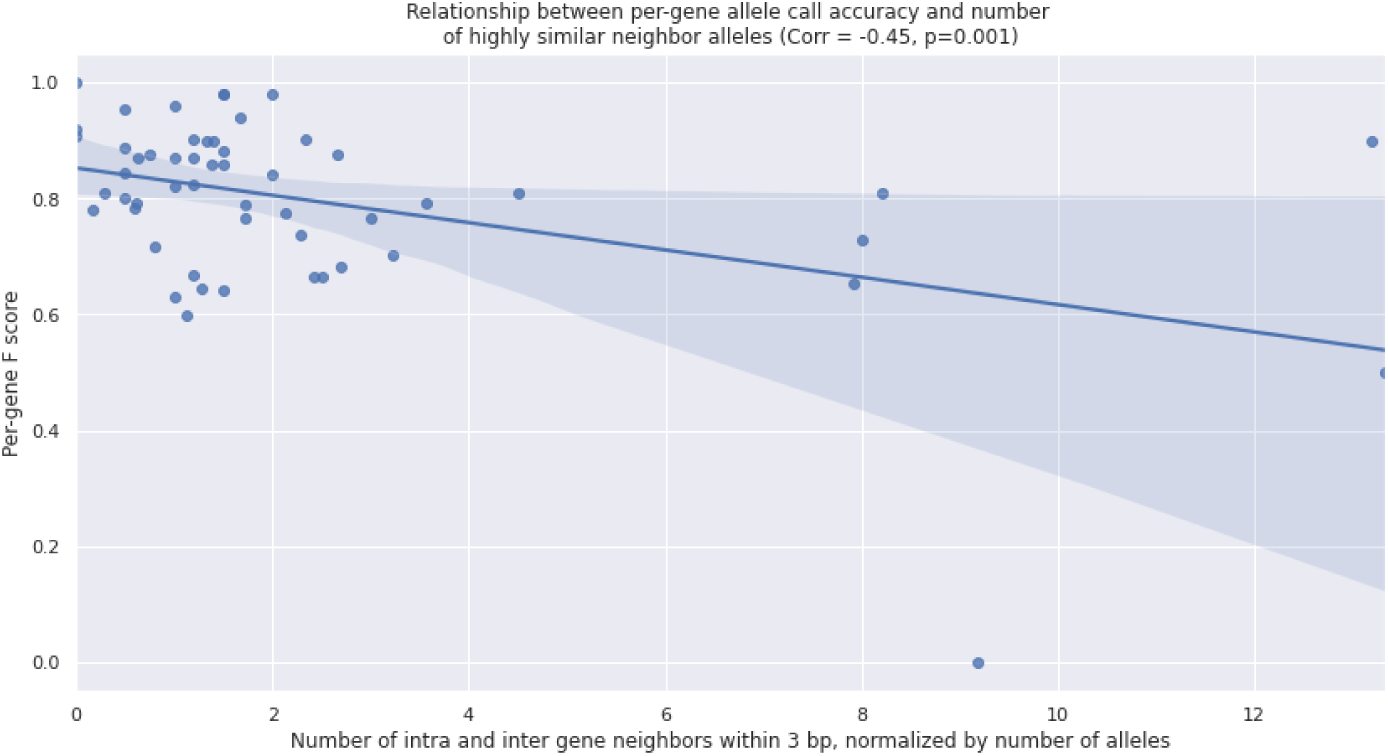
Scatter plot with correlation demonstrating the relationship between ImmunoTyper-SR’s F-score accuracy per gene and the gene’s sequence distinction. The x-axis represents the counts of the number of allele pairs between alleles of a given gene and any other allele in the database that are within 3 bp edit distance. This count is then normalized by the number of alleles known to that gene.

**Figure 4:**
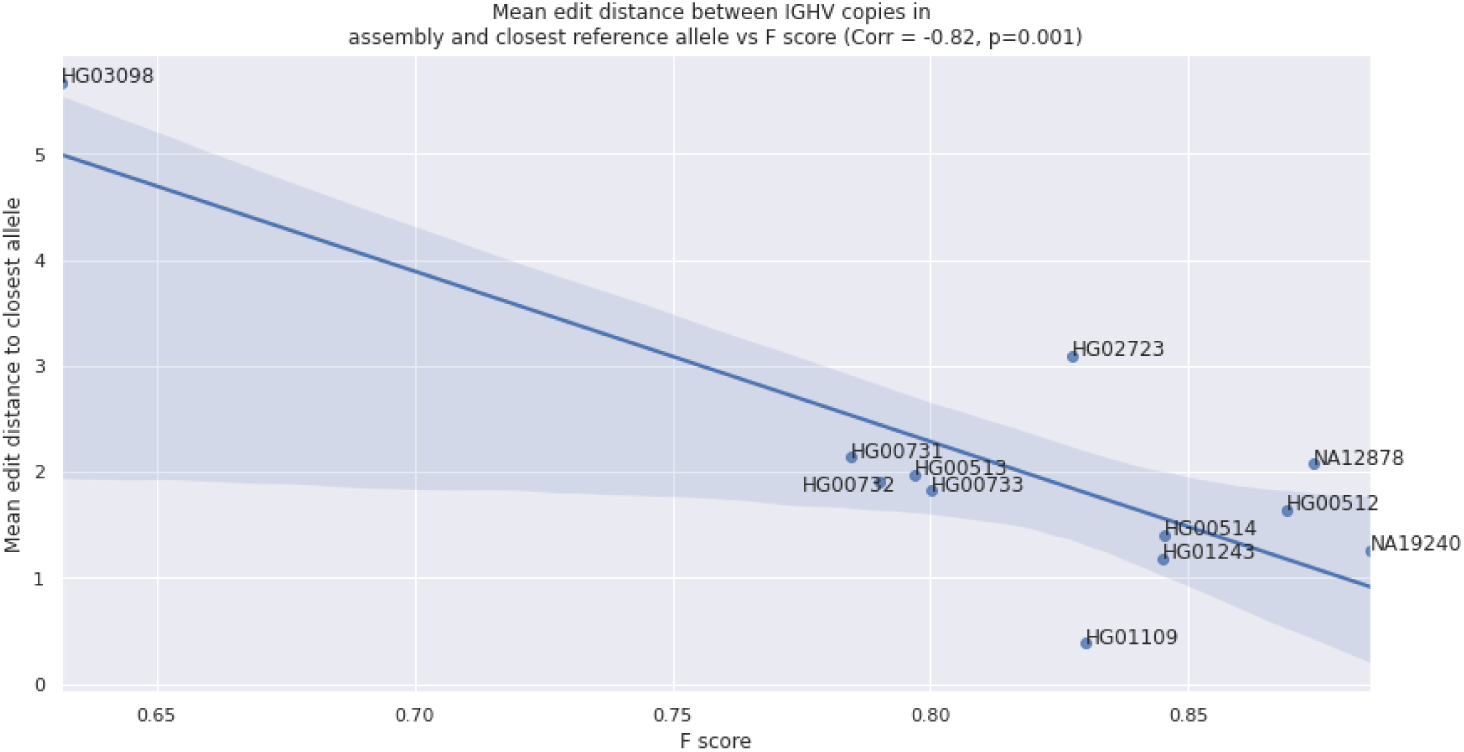
Correlation between the amount of novel variants and genotype call accuracy. The amount of *IGHV* novel variants in a given sample’s *IGH* assemblies is captured as the mean edit distance between each *IGHV* copy and its closest allele in the database. ImmunoTyper-SR genotype call accuracy is represented by the F-score. Corr indicates the Pearson correlation coefficient.

To reduce the impact of this effect, we recalculated allele call accuracy statistics by not penalizing a miscalled (false positive) allele as long as it is within 3 bp edit distance of a ground truth (false negative) allele (see Figure 2). This increases the mean F-score by 6.6%, from 81.5% to 87.9%. The tolerance threshold is chosen to be 3 bp because it is about 1% of the average length of all *IGHV* allele sequences.

#### 3.1.3 ImmunoTyper-SR Reports Genotypes with Higher Granularity and Accuracy than Comparable Methods

We compared ImmunoTyper-SR’s genotype results on the 1kGP samples against the only other published short read *IGHV* genotyping method, as published by Luo et al.[24]. With the exception of at most 11 two-copy genes (out of 56 *IGHV* genes with at least one functional allele), the pipeline reports only gene-level genotypes and CNV calls (see Section 2.6). We implemented the pipeline in-house and compared gene call results in Table 1. We found that ImmunoTyper-SR had significantly higher precision (median 84.7% vs 93.1%), and moderately improved recall (mean 87.4% vs 90.1%).

**Table 1:**
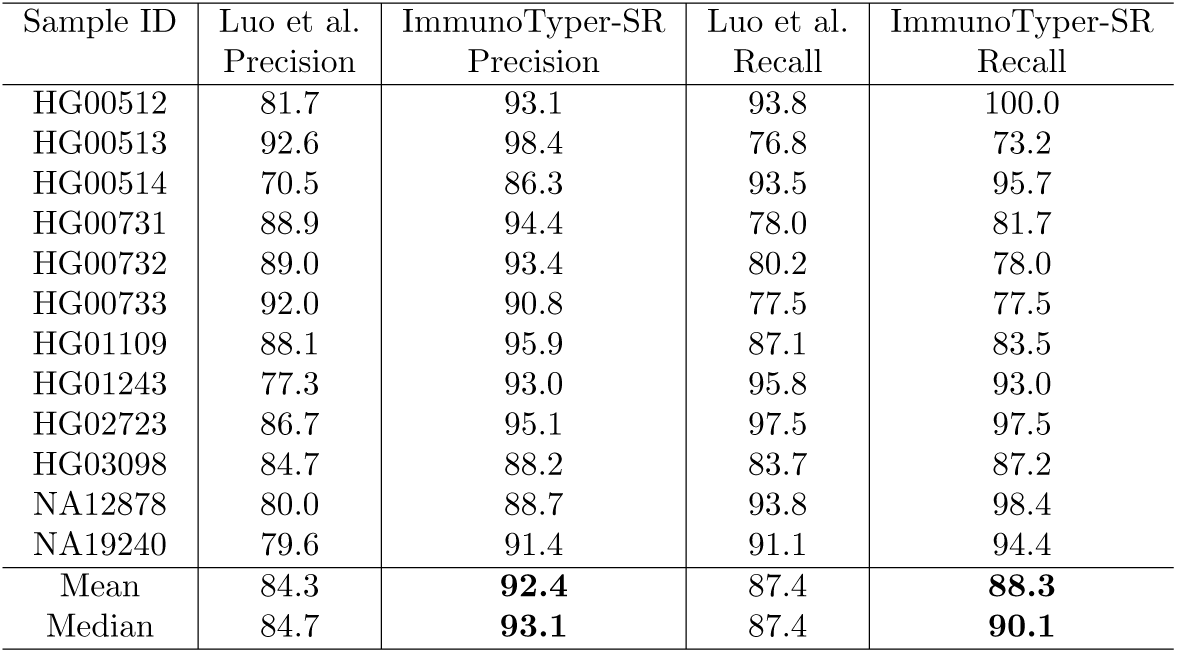
Gene call comparison between Luo et al.[24] pipeline and ImmunoTyper-SR on 1kGP dataset.

| Sample ID | Luo et al.<br>Precision | ImmunoTyper-SR<br>Precision | Luo et al.<br>Recall | ImmunoTyper-SR<br>Recall |
| --- | --- | --- | --- | --- |
| HG00512 | 81.7 | 93.1 | 93.8 | 100.0 |
| HG00513 | 92.6 | 98.4 | 76.8 | 73.2 |
| HG00514 | 70.5 | 86.3 | 93.5 | 95.7 |
| HG00731 | 88.9 | 94.4 | 78.0 | 81.7 |
| HG00732 | 89.0 | 93.4 | 80.2 | 78.0 |
| HG00733 | 92.0 | 90.8 | 77.5 | 77.5 |
| HG01109 | 88.1 | 95.9 | 87.1 | 83.5 |
| HG01243 | 77.3 | 93.0 | 95.8 | 93.0 |
| HG02723 | 86.7 | 95.1 | 97.5 | 97.5 |
| HG03098 | 84.7 | 88.2 | 83.7 | 87.2 |
| NA12878 | 80.0 | 88.7 | 93.8 | 98.4 |
| NA19240 | 79.6 | 91.4 | 91.1 | 94.4 |
| Mean | 84.3 | **92.4** | 87.4 | **88.3** |
| Median | 84.7 | **93.1** | 87.4 | **90.1** |

For allele calls, Luo et al. pipeline calls alleles for only a small proportion of the *IGHV* genes present, an average of 5.75 out of 44 functional genes. For those genes that do have allele calls, ImmunoTyper-SR is significantly more accurate, with mean precision and recall values more than double the Luo et al. results (see Table 2). Note that the 11 genes listed in Section 2.6 are those genes that the authors deemed two copy genes with high confidence, however in many cases not all of those genes were called with two copies and thus not selected for allele calling, despite having two copies in the ground truth.

**Table 2:** Allele call comparison between Luo et al.[24] pipeline and ImmunoTyper-SR. The final column shows the number of ground truth genes in the sample that met the allele-calling criteria of the Luo et al. pipeline (i.e. that the gene is among the 11 least ambiguous genes and its number of copies is predicted to be ≤ 2); their precision and recall values are calculated only on those genes. In contrast, ImmunoTyper-SR calls alleles for all ground truth genes and its precision and recall values are thus calculated on the entire set of genes.

| Sample | Luo et al.<br>Precision | ImmunoTyper-SR<br>Precision | Luo et al.<br>Recall | ImmunoTyper-SR<br>Recall | Proportion of Functional<br>Genes with Allele Calls<br>by Luo et al. |
| --- | --- | --- | --- | --- | --- |
| HG00512 | 50.0 | 94.7 | 50.0 | 100.0 | 9 / 44 |
| HG00513 | 20.0 | 100.0 | 20.0 | 90.0 | 5 / 45 |
| HG00514 | 37.5 | 87.5 | 37.5 | 87.5 | 4 / 34 |
| HG00731 | 56.2 | 100.0 | 56.2 | 93.8 | 8 / 41 |
| HG00732 | 16.7 | 100.0 | 16.7 | 100.0 | 3 / 49 |
| HG00733 | 37.5 | 88.9 | 37.5 | 100.0 | 4 / 48 |
| HG01109 | 41.7 | 100.0 | 45.5 | 100.0 | 6 / 44 |
| HG01243 | 50.0 | 100.0 | 50.0 | 100.0 | 4 / 44 |
| HG02723 | 35.0 | 77.8 | 38.9 | 77.8 | 10 / 44 |
| HG03098 | 50.0 | 76.9 | 50.0 | 71.4 | 7 / 46 |
| NA12878 | 31.2 | 100.0 | 31.2 | 100.0 | 8 / 40 |
| NA19240 | 62.5 | 100.0 | 62.5 | 100.0 | 4 / 49 |
| Mean | 40.7 | **93.8** | 41.3 | **93.4** | 5.75 / 44 |
| Median | 39.6 | **100.0** | 42.2 | **100.0** | 5.5 / 44 |

### 3.2 Validation and Association with anti-Type I IFN Autoantibodies with a Cohort of 585 COVID-19 Patient WGS Data

As part of the NIAID COVID Consortium we analyzed health records and genomic data from a cohort of 585 COVID patients. The dataset includes various health information for each patient, as well as whole genome sequences performed at The American Genome Center, USUHS, and results from anti-type I IFN autoantibody (aAb) assays.

#### 3.2.1 Genotype Validation Through Doubly-Sequenced Concordance

To further validate ImmunoTyper-SR on non-LCL sourced WGS samples, we compared genotype calls for nine individuals who had been independently sequenced twice as part of the NIAID cohort. The WGS were generated at the same sequencing center, which reduces the effect of systematic sequencing bias. We compared the complete set of allele calls and CNVs between the matched WGS. The mean weighted Jaccard similarity coefficient across all nine doubly-sequenced samples was 0.696. If the CNV calls are ignored, the mean score increases to 0.923, indicating many of the miscalls are due to CNVs, rather than calling the wrong allele.

#### 3.2.2 Association with anti-Type I IFN Autoantibodies

Removing samples that did not have WGS or aAb assay results produced 542 samples, of which 32 tested positive for the presence of anti-type I IFN aAb. The association, effect size, and *p*-values are provided in Table 4 for the top most significant alleles. It is important to note that three out of four of the top most significant alleles are rare alleles, present in at most 10 individuals. In addition, this analysis suffers from a very low number of Type I IFN aAb cases present in this dataset. While two of the selected alleles have relatively low *p*-values, they remain higher than 0.01, partially due to their rarity in the cohort.

**Table 3:** Distribution of Jaccard similarity coefficients for doubly-sequenced samples in the NIAID cohort: (column 2) no error tolerance in sequence composition or copy number of called alleles; (column 3) no error tolerance in copy number but an edit distance of ≤ 3 tolerated on called alleles; (column 4) no error tolerance in sequence composition of the alleles with copy number differences ignored.

| Subject ID | Copy number-sensitive | Copy number-sensitive<br>3 bp allowance | Copy number-insensitive |
| --- | --- | --- | --- |
| CV01 | 0.610 | 0.664 | 0.902 |
| CV02 | 0.752 | 0.777 | 0.913 |
| CV03 | 0.683 | 0.702 | 0.943 |
| CV04 | 0.689 | 0.719 | 0.942 |
| CV05 | 0.784 | 0.800 | 0.956 |
| CV06 | 0.708 | 0.744 | 0.927 |
| CV07 | 0.704 | 0.765 | 0.903 |
| CV08 | 0.680 | 0.719 | 0.901 |
| CV09 | 0.656 | 0.678 | 0.925 |
| Mean | 0.696 | 0.730 | 0.923 |
| Median | 0.693 | 0.724 | 0.924 |

**Table 4:** Summary statistics of *IGHV* alleles that are associated with Type I IFN aAb in the 542 samples from the NIAID cohort that have been tested. Top row: the proportion of the samples with each allele. Second row: the number of samples with the allele as well as the Type I IFN aAb. Third row: Number of samples with the allele but not the IFN aAb. Bottom two rows: Association between the allele and the Type I IFN aAb.

| Allele | *IGHV*1-46*Novel-168 | *IGHV*5-51*Novel-501 | *IGHV*3-49*04 | *IGHV*3-9*Novel-205 |
| --- | --- | --- | --- | --- |
| Samples with genotype | 3 / 542 | 10 / 542 | 237 / 542 | 9 / 542 |
| Samples with IFN aAb | 1 | 2 | 10 | 2 |
| Samples without IFN aAb | 2 | 8 | 227 | 7 |
| Effect probability | 37.5 | 24.6 | 3.7 | 26.6 |
| P-value | 0.037 | 0.018 | 0.11 | 0.013 |

## 4 Discussion and future work

ImmunoTyper-SR successfully raises the bar for short read-based *IGHV* germline genotyping. While the tool has demonstrated the greatest accuracy of published Illumina-based tools to date, it is important to note that there is still room for improvement, particularly when distinguishing highly-similar alleles.

The challenges presented by novel alleles are another opportunity for improving *IGHV* genotype calls. ImmunoTyper-SR clearly performs best when a sample’s *IGHV* alleles minimally diverge from the known allele database. As the *IGHV* allele database is updated and improved, genotype call accuracy will likely increase. Ongoing projects such as the OGRDB[37], a curated database of germline *IGHV* alleles inferred from AIRR-seq data, as well as those being curated from high-quality long read assemblies, may help improve WGS short read-based germline *IGHV* calling. In addition, our future work includes expanding ImmunoTyper-SR to call novel alleles and variants.

This paper represents the first investigation into germline *IGHV* genotype-phenotype association using short read WGS data. Despite there being a hypothetical biological mechanism for the association between *IGHV* genotype and anti-type I IFN autoantibodies, the dataset used is very limited for this application. The low number of autoantibody-positive patients and rarity of the associated alleles severely reduces statistical power resulting in a low-confidence statistical analysis. As a result, we must judge the association results as inconclusive, despite finding two associated alleles with relatively low *p*-values. However, our future work includes repeating this analysis on a cohort of *>* 1200 COVID-19 patient WGS data, which we believe will improve our confidence.

While our investigation of association was inconclusive, there are many other potential association targets for future work. It is important to acknowledge a possibility that disease effects of germline *IGHV* variants can be overcome by somatic hypermutation during antibody repertoire proliferation. However, the (currently limited) body of evidence demonstrating *IGHV* variant effects on phenotype suggests there are many *IGHV* alleles that have a significant effect on a variety of diseases. In the near future, comprehensively investigating these associations is likely only possible using short read WGS data; there are numerous WGS datasets in existence which have remained untouched with respect to *IGHV*, and alternative data types have additional costs and scaling challenges.

It is possible that using alleles as the explanatory variables in phenotype association is less biologically relevant than single nucleotide variants. If a single nucleotide variant is the root driver of a phenotype effect, perhaps by affecting development of a given protective or detrimental antibody, then using variants as an explanatory variable would be more suitable. However, if the effect is driven by the combination of variants, using alleles may be more suitable. Given a sufficient sized dataset, joint effects, either of variants or alleles, could also be investigated. It is also possible that other variant types, such as structural variants or CNVs, could have an effect on phenotype. Our future work includes improving ImmunoTyper-SR to provide variant calls, which will improve the ability to investigate phenotype and disease associations.

There is significant opportunity for future studies that combine WGS with complete BCR repertoire data. The immediate application would be to validate WGS and inference-based germline *IGHV* genotype calls using orthogonal datasets. Unfortunately we were unable to find any such paired datasets for this study.

There are also open questions regarding the direct effects of germline *IGHV* genotype on the BCR repertoire. This effect has been investigated with repertoire data alone, however inference-based germline genotype calls are impacted by temporal effects of repertoire sampling, and as a result may be incomplete. By combining germline sequence data with repertoire data, it will be possible to create a complete picture of the interaction between germline *IGHV* genotype and the BCR repertoire.

As the first short read WGS-based *IGHV* genotyping tool with allele-level granularity, ImmunoTyper-SR opens the door to applying the power of the largest WGS datasets available to uncover the mysteries of one of the least understood loci in the human genome.

## Code availability

Source code will be available through GitHub.

## Acknowledgements

This work was supported by funding from the the Intramural Research Programs of the National Cancer Institute, and the National Institute of Allergy and Infectious Diseases, National Institutes of Health. Ethics approval was obtained from the University of Milano-Bicocca School of Medicine, San Gerardo Hospital, Monza–Ethics Committee of the National Institute of Infectious Diseases Lazzaro Spallanzani (84/2020) (Italy), and the Comitato Etico Provinciale (NP 4000–Studio CORONAlab). STORM-Health care workers were enrolled in the STudio OsseRvazionale sullo screeningdei lavoratori ospedalieri per COVID-19 (STORM-HCW) study, with approval from the local institutional review board (IRB) obtained on 18 June 2020. The WGS data has been deposited in dbGaP (accession: `phs002245`^4^).

## Disclaimer

The content of this publication does not necessarily reflect the views or policies of the Department of Health and Human Services, nor does mention of trade names, commercial products, or organizations imply endorsement by the U.S. Government.

## Footnotes

^1^https://www.niaid.nih.gov/research/host-genetics-severe-covid-19-infection

2 A complementary international effort, the COVID-19 host genetics initiative (https://www.covid19hg.org) has similar goals.

3 i.e., the edit distance between such a pair of alleles is no more than 2% of the length of the shorter allele

4 https://www.ncbi.nlm.nih.gov/projects/gap/cgi-bin/study.cgi?study_id=phs002245.v1.p1

